# Regulator of Chromosome Condensation (RCC1) a novel therapeutic target in pancreatic ductal adenocarcinoma drives tumor progression via the c-Myc-RCC1-Ran axis

**DOI:** 10.1101/2023.12.18.572102

**Authors:** Sahar F. Bannoura, Amro Aboukameel, Husain Yar Khan, Md. Hafiz Uddin, Hyejeong Jang, Eliza Beal, Amalraj Thangasamy, Seongho Kim, Kay Uwe Wagner, Ramzi Mohammad, Mohammed Najeeb Al-Hallak, Boris C. Pasche, Asfar S. Azmi

## Abstract

Pancreatic ductal adenocarcinoma (PDAC) is a highly lethal malignancy with limited therapeutic options. Here we for the first time evaluated the role of regulator of chromosome condensation 1 (RCC1) in PDAC subsistence and drug resistance. RCC1 expression was found to be elevated in PDAC tissues in comparison with normal pancreatic tissues and was linked to poor prognosis. RCC1 silencing in a panel of PDAC cells by RNA interference and CRISPR-Cas9 resulted in reduced cellular proliferation in 2D and 3D cultures. RCC1 KD reduced migratory and clonogenic ability, enhanced apoptosis, and altered cell cycle distribution in human PDAC cells as well as cells isolated from the LSL-Kras^G12D/+;^LSL-Trp53^R172H/+^;Pdx1-Cre (KPC) mouse tumors. Subcutaneous cell-derived xenografts show significantly attenuated growth of RCC1 KO tumors. Mechanistically, RCC1 knockdown resulted in disruption of subcellular Ran distribution indicating that stable nuclear Ran localization is critical for PDAC proliferation. Nuclear and cytosolic proteomic analysis revealed altered subcellular proteome in RCC1 KD KPC-tumor-derived cells. Altered cytoplasmic protein pathways include several metabolic pathways and PI3K-Akt signaling pathway. Pathways enriched in altered nuclear proteins include cell cycle, mitosis, and RNA regulation. RNA sequencing of RCC1 KO cells showed widespread transcriptional alterations. Upstream of RCC1, c-Myc activates the RCC1-Ran axis, and RCC1 KO enhances the sensitivity of PDAC cells to c-Myc inhibitors. Finally, RCC1 knockdown resulted in the sensitization of PDAC cells to Gemcitabine. Our results indicate that RCC1 is a potential therapeutic target in PDAC that warrants further clinical investigations.

## INTRODUCTION

Pancreatic ductal adenocarcinoma (PDAC) is an unmet clinical problem requiring urgent attention. In the United States, the 5-year relative survival rate is alarmingly low at 12%^1^, underscoring the need for more effective therapies. Currently, chemotherapy is the mainstay of treatment for the majority of PDAC patients, but its efficacy remains limited. KRAS mutations have been identified as drivers of PDAC in more than 90% of cases. However, the long-awaited advent of KRAS-targeted therapies, including the G12C and G12D mutant KRAS inhibitors, remain elusive for PDAC patients, representing a significant gap in targeted therapeutics.

The Regulator of Chromosome Condensation 1 (RCC1) is a guanine exchange factor (GEF) for the Ras-related nuclear protein (Ran). It promotes the activation of Ran through the exchange of Ran-bound GDP for GTP to ensure proper nucleocytoplasmic transport, mitosis, and nuclear envelope formation. RCC1 is a chromatin-bound protein, that binds nucleosomes and double stranded DNA ^2,3^, where it generates high local concentrations of Ran-GTP. High levels of Ran-GTP around chromatin are critical for microtubule aster formation and the proper assembly of the mitotic spindle ^2,4,5^. At the end of mitosis, RCC1 plays a role in nuclear pore complex and nuclear envelope reassembly. Notably, RCC1 obtained its name because of its involvement in the regulation of the onset of chromosome condensation in S phase of the cell cycle ^6^.

Ran is a small GTPase that acts as the primary regulator of nucleocytoplasmic transport, governing the import and export of proteins and RNAs in and out of the nucleus across the nuclear membrane. Ran cycles between an active GTP-bound state in the nucleus and an inactive GDP-bound state in the cytosol as regulated by nuclear RCC1 GEF activity, and cytosolic Ran GTPase activating protein (RanGAP1) activity, respectively. The resulting gradient of Ran-GTP is important for maintaining the correct direction of molecular cargo transport. Active nucleocytoplasmic transport is required for proliferative cells to propagate signals from outside the cell into the nucleus, and to support proliferative activities. Highly proliferative cells require a steep Ran-GTP gradient where cancer cells have a steeper gradient than normal cells^7^.

RCC1 has been implicated in DNA damage response (DDR) by ensuring the proper localization of DDR factors and their recruitment to sites of DNA damage. ATM and ATR kinases are imported into the nucleus in a Ran-dependent manner. Moreover, RCC1 overexpression gives normal cells the ability to evade the DNA-damage checkpoint and progress through the cell cycle^8^. This indicates that high RCC1 levels may result in tolerance to DNA damage, and replication of cells with unrepaired DNA. On the other hand, a deficiency in RCC1 may result in an improper response to DNA damage. These processes are of critical importance for tumorigenesis, as well as response to cancer therapeutics.

We were the first group to define the role of RCC1 in PDAC ^9^. Using CARIS database, DNA and RNA sequencing was performed on 5,071 PDAC tissues. High RCC1 expression was associated with worse overall survival (p<0.00001; HR 1.522) and worse overall survival from the start of chemotherapy (p<0.0001; HR 1.494) in PDAC. Here, we characterized the role of RCC1 in PDAC as a potential therapeutic target. We found that RCC1 is overexpressed in PDAC, which drives tumor growth, and proteome-wide perturbations. RCC1 silencing in PDAC cells resulted in sensitization of cells to chemotherapy treatment.

## RESULTS

### RCC1 is Elevated in PDAC and is Associated with Poor Prognosis

We sought to characterize RCC1 expression in PDAC tumors, and how it relates to survival. We analyzed publicly available data from the TCGA, GTEx, and CPTAC databases. Normal and tumor RNA-seq data was analyzed using the Xena platform (UCSC) (Figure 1A). Comparison of expression levels revealed that RCC1 expression is elevated in PDAC in comparison to normal tissues in a statistically significant manner (p=1.009e-9; Figure 1A). RCC1 expression in formalin fixed paraffin embedded (FFPE) patient tissues compared to normal tissues showed a similar trend (Figure S1A). Additionally, comparison of protein expression revealed higher RCC1 levels in PDAC compared to normal tissues (p=0.025; Figure 1B). Higher RCC1 expression is associated with a worse five-year overall survival rate in TCGA PDAC patients (p=0.0048; Figure 1C). Under a rapid autopsy program (RAP), we were able to collect tissues from a patient with metastatic PDAC. IHC analysis of RAP tissues confirmed upregulation of RCC1 expression in PDAC tissues compared to adjacent normal pancreatic tissues (Figure 1D). We also investigated RCC1 levels in a tissue microarray (TMA) containing normal pancreatic and malignant PDAC tissues of different grades. This analysis showed increased intensity of staining in the nuclei of adenocarcinoma cells in the malignant tissues (Figure 1E). Additionally, cells within the tumor stroma stained positively for RCC1, whereas normal stromal cells did not show such staining (Figure 1E). This increased staining intensity was also higher in grade 3 tumors, compared to grade 1 and 2 (Figure 1F), and higher in stage III patients compared to stages I and II (Figure S1B), suggesting an increase in RCC1 expression as the tumor progressed. Next, we analyzed RCC1 expression in a panel of PDAC cell lines and found significant RCC1 expression of various degrees in different cell lines, including cells with oncogenic KRAS G12C and G12D mutations, as well as KRAS WT cells (Figure 1G). Overall, these data show that there is increased expression of RCC1 in PDAC, and it may be correlated with disease progression and worse prognosis.

**Figure 1.**
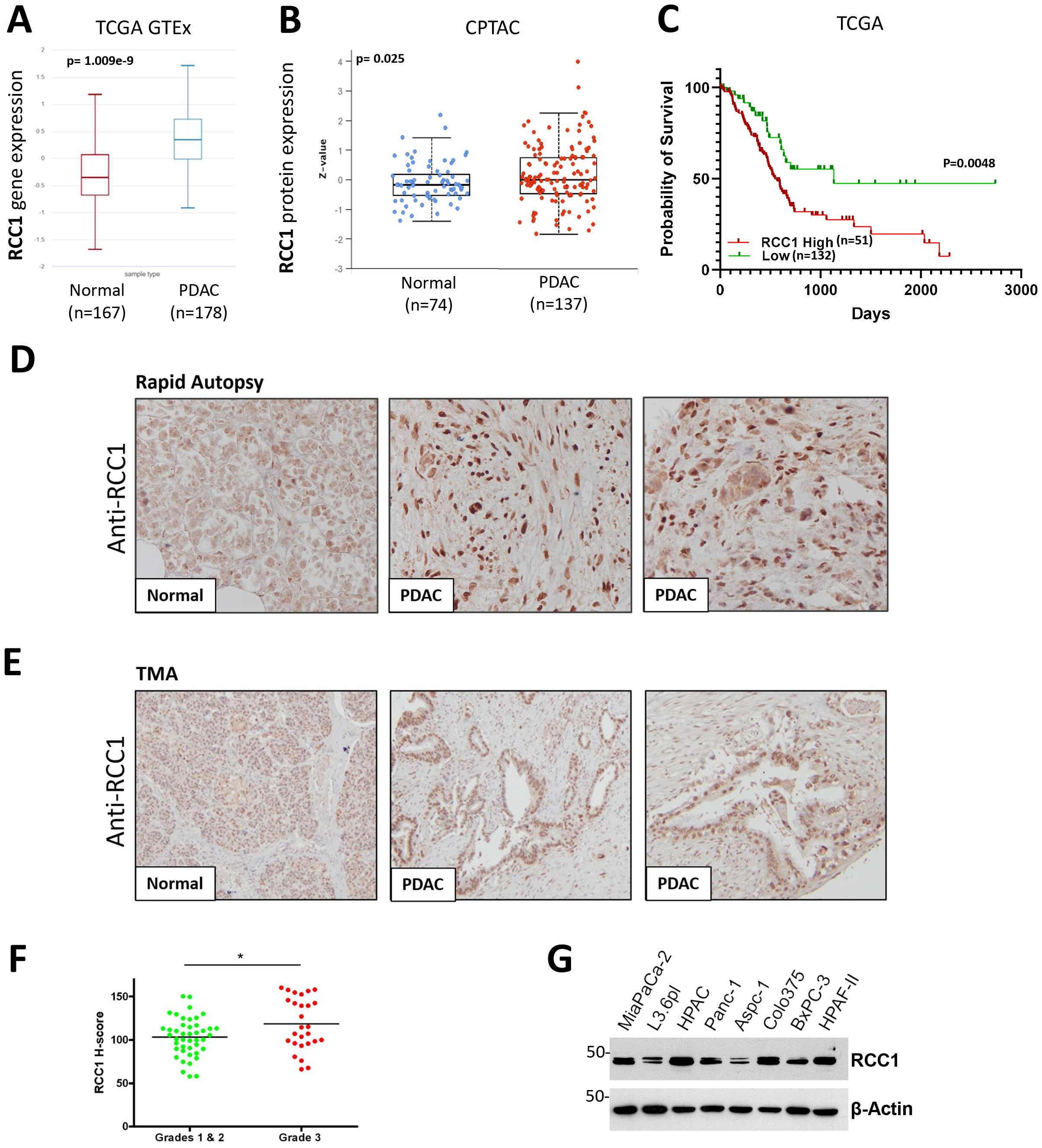
*RCC1* expression is elevated and is associated with poorer prognosis in PDAC. (A) Box plot shows *RCC1* RNA expression in pancreatic normal and tumor tissues (n=350). Reads were realigned and reanalyzed using the same pipeline to eliminate any bias or batch effect. (B) Box plot showing RCC1 protein expression levels in normal pancreatic or PDAC tissues from the CPTAC database. (C) Kaplan Meier curve depicting overall survival in TCGA patients according to *RCC1* status (n=183). Samples were obtained from patients who underwent pancreatectomy or Whipple procedure prior to therapy. Stage distribution of the patients: stage I: 10%; stage II: 83%; stage III: 3%; stage IV: 4%. (D) Micrographs of PDAC and adjacent normal tissues obtained through a rapid autopsy program stained for RCC1 by immunohistochemistry. (E)Micrographs of normal or PDAC tissues from a tissue microarray (TMA) stained as in (D). (F)Dot plot of TMA RCC1 H-scores according to Grade. (G) Western blot of RCC1 in a panel of PDAC cell lines.

### RCC1 depletion in PDAC cells inhibits tumor properties

To determine the functional role of RCC1 in PDAC, we knocked down the expression of RCC1 using RNA interference (siRNA) in MIA PaCa2, BxPC3, AsPC1 and PanC1 cell line models of PDAC (Figure 2A). These cell lines differ in their characteristics, including morphology, mutational status, and growth rate. RCC1 inhibition resulted in significantly decreased cell viability across the four different cell lines (Figure 2B), decreased cellular migration (Figure 2C), and reduced capacity for colony formation (Figure 2D). Additionally, we utilized a PDAC cell model derived from a pancreatic tumor of the **K**ras(G12D/^+^); LSL-Tr**p**53(R172H/^+^); Pdx1-**C**re model (KPC), termed KCI313 (Figure S1C). The KPC mouse model is a well-studied and one of the most utilized GEMMs of PDAC since it develops pancreatic cancer that is reminiscent of human disease in genetic, histological, and clinical aspects^10^. We confirmed that the KCI313 were positive for Kras G12D and Trp53 mutations (data not shown). Murine Rcc1 siRNA-mediated silencing in these cells resulted in decreased cell viability in two-dimensional and in three-dimensional spheroid models (Figure 2E). Additionally, Rcc1 silencing resulted in increased cellular apoptosis (Figure S1D) and altered cell cycle distribution showing cells arrested in the S and G2M phases (Figure S1E). On the other hand, overexpression of Rcc1 in KCI313 cells resulted in increased cellular proliferation (Figure S1F). To further characterize the role of RCC1 in PDAC cells, we utilized CRISPR-Cas9 targeting to knock out RCC1 in HPAF-II and MIA PaCa2 cells (Figure 3A). RCC1 knockout significantly altered the properties of PDAC cells (Figure 3B-G), indicating that RCC1 is critical for PDAC cells. Depletion of RCC1 resulted in decreased viability of HPAF-II and MIA PaCa2 knockout cells. Clones from both cell lines were grown to isolate cells showing efficient RCC1 knockout (Figure S2A-C, Figure 3E). RCC1 knockout cells also showed reduced clonogenic capacity (Figure 3C), as well as reduced spheroid growth (Figure 3D). Clonally isolated cells also showed reduced clonogenic capacity (Figure 3E, F). RCC1 depleted cells also demonstrated aberrant cell cycle progression, with cells being arrested in the S and G2/M phases indicating replicative stress, and reduced capacity at undergoing mitosis (Figure 3G, Figure S2D, E).

**Figure 2.**
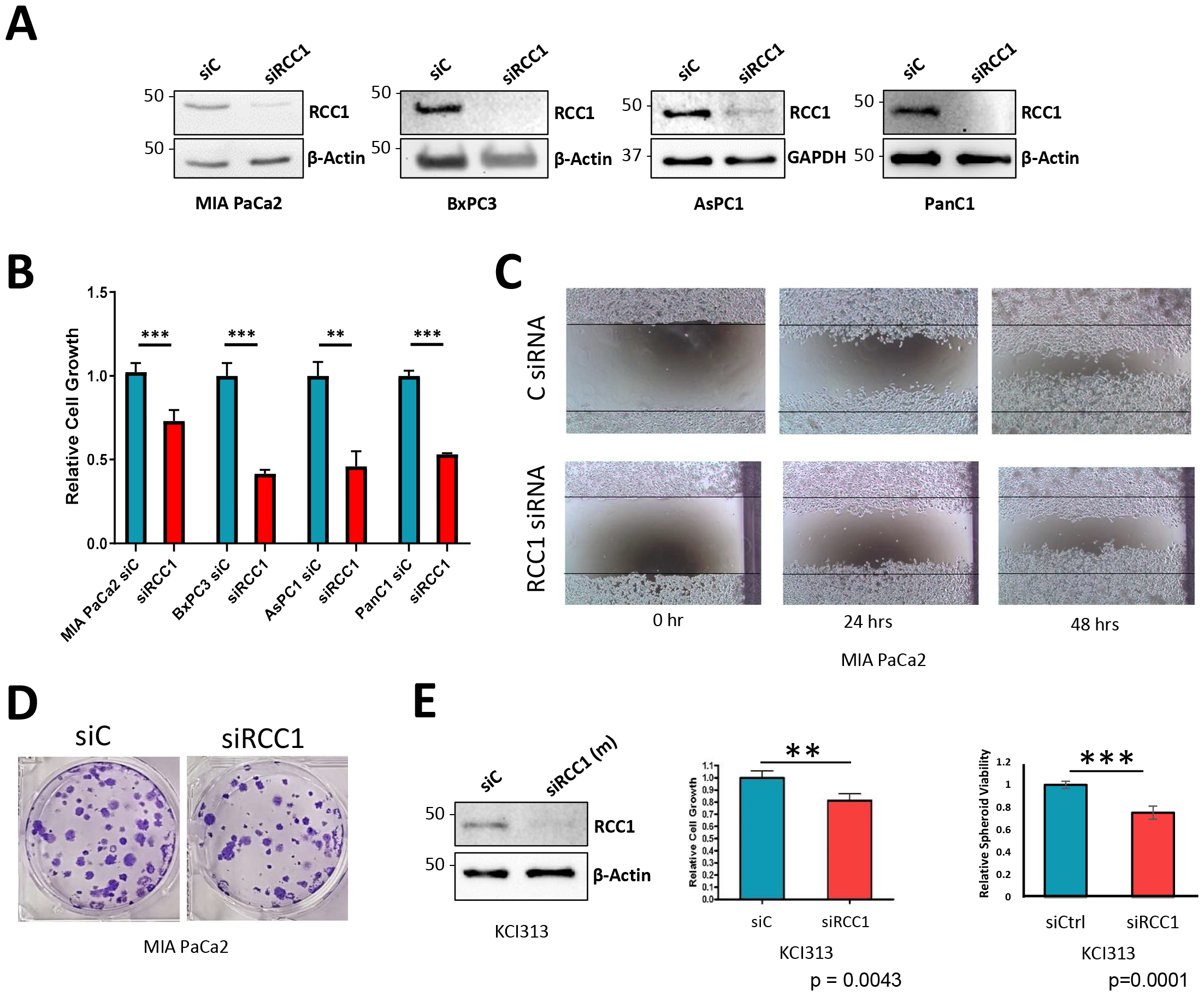
*RCC1* silencing inhibits the proliferation of PDAC cells. (A) Western blot showing efficiency of siRNA mediated *RCC1* silencing at 72 hrs. (B) Bar graph showing reduced cell viability in *RCC1* KD PDAC cells (72 hrs). (C) Scratch wound healing assay showing reduced cellular migration in MIA PaCa2 upon *RCC1* KD. (D) Colony formation assay showing reduced colony number and size in *RCC1* KD MIA PaCa2 cells. (E) KPC derived KCI313 cells were transfected with mouse specific *RCC1* siRNA or a non-targeting control. Western blot showing knockdown efficiency, and bar graphs showing reduced cell viability 72 hrs after knockdown (left), and reduced spheroid size in knockdown cells (right).

**Figure 3.**
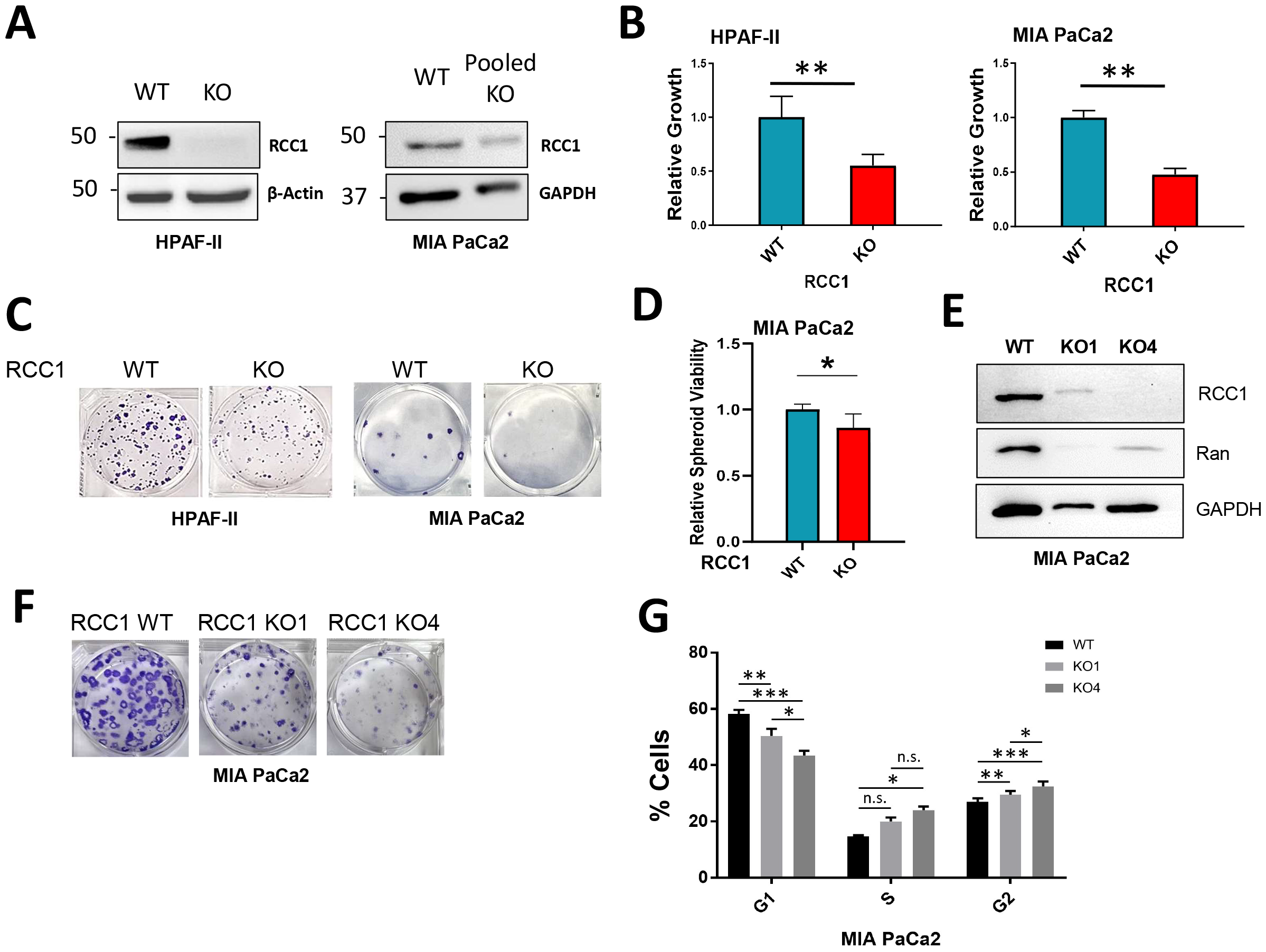
*RCC1* knockout using CRISPR-Cas9 attenuates cellular proliferation and alters cell cycle. (A) Western blot showing efficiency of *RCC1* KO in HPAF-II and MIA PaCa2 PDAC cell lines. (B) Bar graph showing reduced growth of *RCC1* KO cells. (C) Clonogenic assay demonstrating reduced colony number and size in *RCC1* KO cells. (D) Bar graph depicting smaller spheroids in MIA PaCa2 *RCC1* KO cells. (E) Western blot showing RCC1 and Ran levels in MIA PaCa2 WT and two isolated *RCC1* KO clones termed KO1 and KO4. (F) Reduced colony formation and size in MIA PaCa2 *RCC1* KO1 and KO4 clones. (G) Bar graph depicting alteration in cell cycle progression in *RCC1* KO1 and KO4 MIA PaCa2 cells.

To study the role of RCC1 in tumor growth *in vivo*, we compared the growth patterns of parental HPAF-II cells, and MIA PaCa2 cells, with HPAF-II and MIA PaCa2 cells with RCC1 KO, respectively using bilateral subcutaneous xenograft implantation (Figure 4A, B). RCC1 depletion in HPAF-II cells resulted in restricted tumor growth in two independent experiments (Figure 4A, figure S3A. Harvested tumors showed a significant difference in size and weight (Figure 4A, figure S3A), and the persistence of RCC1 knockout in the tumors was confirmed (Figure S3B). MIA PaCa2-derived tumors also showed differential growth based on RCC1 status, where tumor growth was significantly diminished in RCC1 knockout tumors (Figure 4B). These results indicate that RCC1 is important for proliferation and its silencing results in tumor growth inhibition. Immunohistochemical staining showed lower nuclear Ran expression in RCC1 knockout HPAF-II (Figure 4C). This suggests that RCC1 is acting upstream of Ran to regulate its expression and localization in PDAC, thereby altering growth patterns.

**Figure 4.**
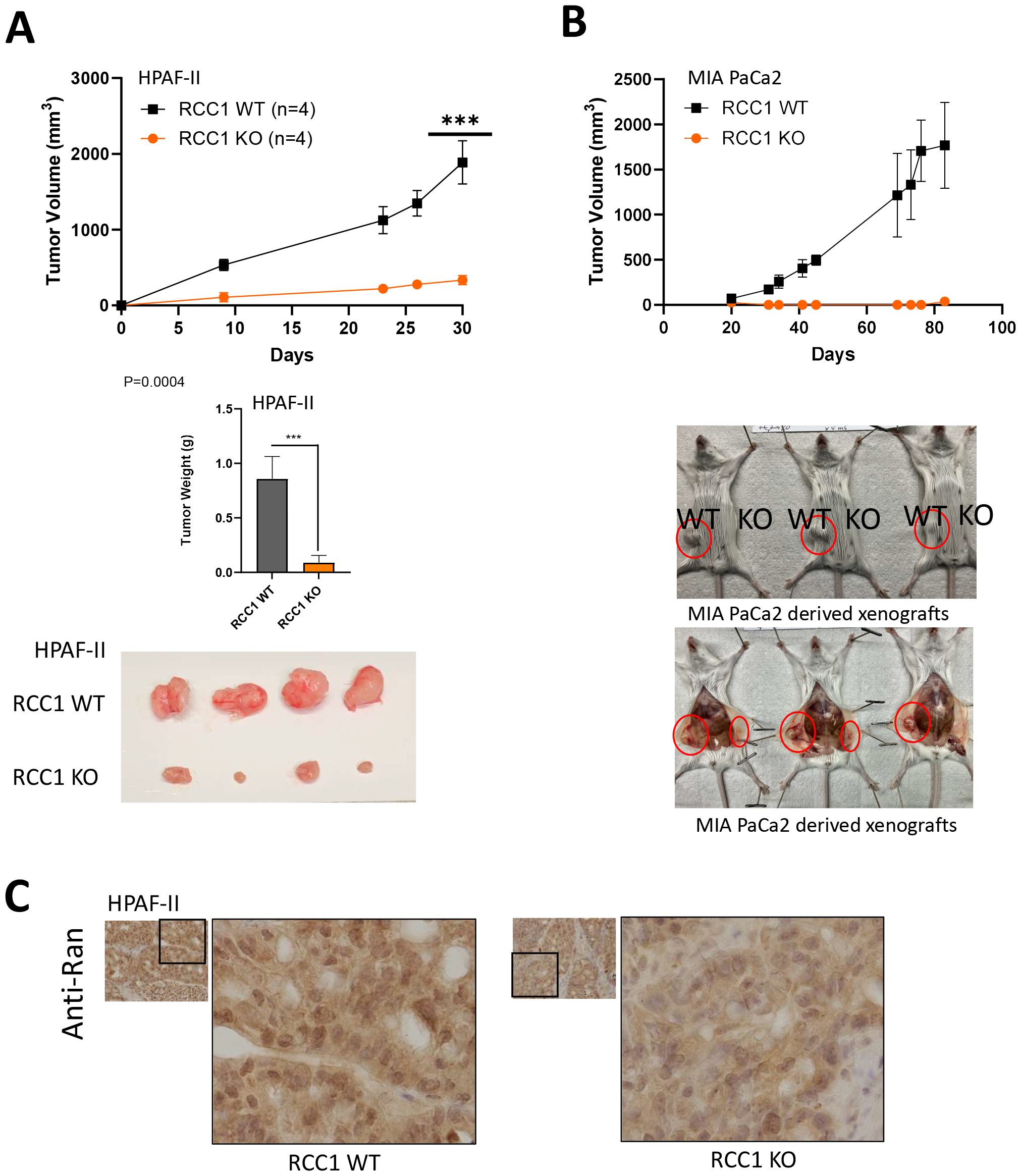
*RCC1* KO restricts PDAC tumor growth in vivo. **(A)** An equal number of HPAF-II WT and HPAF-II *RCC1* CRISPR/Cas9 knockout cells (1 × 10^6^) were bilaterally injected subcutaneously in ICR-SCID mice. Growth was monitored over 1 month (*** p<0.001). Bar graph shows difference in tumor weights. Picture of harvested tumors is shown. **(B)** same as (A) for MIA PaCa2 followed over 3 months. Pictures of tumors in mice after euthanasia is shown. (C) Immunohistochemical staining of HPAF-II derived xenografts showing suppression of Ran upon RCC1 *KO*.

### RCC1 silencing induces proteome wide alterations in protein localization

We observed that Ran expression decreased in RCC1 KO clones of MIA PaCa2 (Figure 3E, Figure S2B), and decreased in HPAF-II derived xenografts (Figure 4C). These observations indicated that RCC1 is likely regulating Ran expression and localization in PDAC, warranting further investigation into the significance of Ran in PDAC, and how it relates to RCC1 function. Based on these observations and functions of RCC1 and Ran in nucleocytoplasmic transport), we employed a proteomics analysis of nuclear and cytoplasmic proteins in KCI313 cells. We found a proteome wide perturbation of protein localization in the nuclear and cytosolic compartments of the cells (Figure 5A, B; Figure S4). In the cytoplasm, 230 proteins were downregulated, and 271 proteins were upregulated. Likewise in the nucleus, 114 proteins were downregulated, and 166 proteins were upregulated. KEGG pathway analysis of the proteins that are altered in the nucleus and cytoplasm revealed multiple pathways that are impacted by protein redistribution (Figure 5B). These pathways included metabolic biosynthesis pathways of amino acids, tRNA and unsaturated fatty acids. This indicates that RCC1 may be mediating a metabolic rewiring within PDAC cells. Cell cycle and RNA transport pathways were also enriched, which validates our model since these are canonical RCC1 functions ^11^.

**Figure 5.**
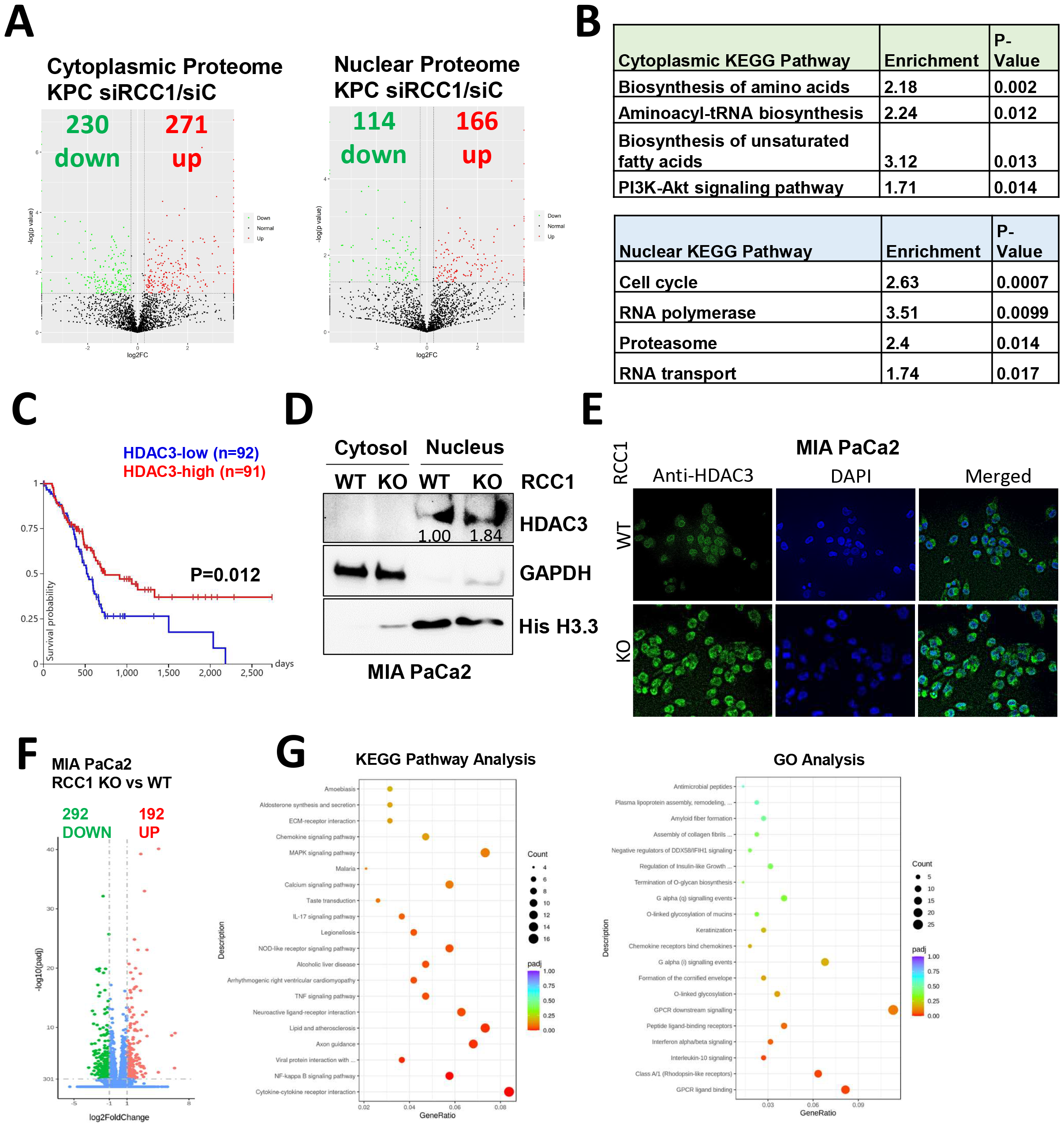
*RCC1* silencing alters the proteome and transcriptome of PDAC cells. **(A)** Volcano plots of differentially expressed proteins in nuclear and cytoplasmic fractions of *RCC1* silenced KPC derived cells. **(B)** Selected significantly enriched pathways in the proteomics dataset. **(C)** Kaplan Meier plot demonstrating difference in overall survival of PDAC patients in the TCGA based on *HDAC3* expression. **(D)** Nuclear cytoplasmic fractionation showing elevated nuclear HDAC3 in *RCC1* KO cells. **(E)** Immunofluorescence staining of HDAC3 in MIA PaCa2 *RCC1* WT and KO cells. **(F)** Volcano plot of differentially expressed genes (DEG) in MIA PaCa2 *RCC1* KO cells. Downregulated genes = 292. Upregulated genes = 192. **(G)** KEGG pathway and gene ontology analysis of DEGs in *RCC1* KO MIA PaCa2 cells.

HDAC3 plays a key role in epigenetic regulation of gene expression, cell cycle progression, and development, and has been implicated in pancreatic cancer ^12-14^. Proteomics analysis revealed HDAC3 upregulation in the nuclear fraction. TCGA overall survival data shows that patients with high HDAC3 have a more favorable prognosis compared to low HDAC3 (Figure 5C). Validation in a human PDAC cell line, MIA PaCa2, confirmed an almost 2-fold increase in HDAC3 in the KO cells using western blotting and immunofluorescence (Figure 7 D, E). As a transcriptional repressor, it is possible that HDAC3 may be playing a compensatory role upon RCC1 silencing, which makes it a potential co-target with RCC1.

To investigate transcriptional alterations upon RCC1 silencing, we conducted whole exome RNA sequencing comparing MIA PaCa2 cells with RCC1 WT or KO. This revealed the downregulation of 292 genes, and the upregulation of 192 genes in the RCC1 KO cells compared to WT (Figure 5F). In order to gain insight into the functions of the differentially expressed genes, we performed KEGG pathway and Gene Ontology (GO) analyses (Figure 5G). The top altered pathways were cytokine-cytokine receptor interaction, and NF-kappa B signaling pathway. Top enriched GO terms included GPCR ligand binding, and GPCR downstream signaling.

### c-Myc induces RCC1 to regulate Ran expression and localization in PDAC

Thus far, our data show that RCC1 is acting upstream of Ran, and modulation of the RCC1-Ran axis is important for PDAC. We sought to delineate the impact of RCC1 modulation on Ran downstream. First, we analyzed PDAC patient survival in the TCGA database based on RAN expression. Patients were divided into quartiles according to RAN expression levels. We found that patients in the RAN high quartile have worse overall survival (Figure 6A left), and worse progression free survival (Figure 6A right) compared to the patients in the low RAN expression quartile. To investigate how Ran localization is affected by RCC1 depletion, we fractionated the cells into their cytoplasmic and nuclear protein compartments, which revealed that Ran distribution is altered in RCC1 knockout cells, where nuclear Ran concentrations were depleted in the RCC1 knockout cells (Figure B). These results indicate that RCC1 activity is indispensable for maintaining the steady state Ran distribution, as well as its nuclear localization for proper functioning.

**Figure 6.**
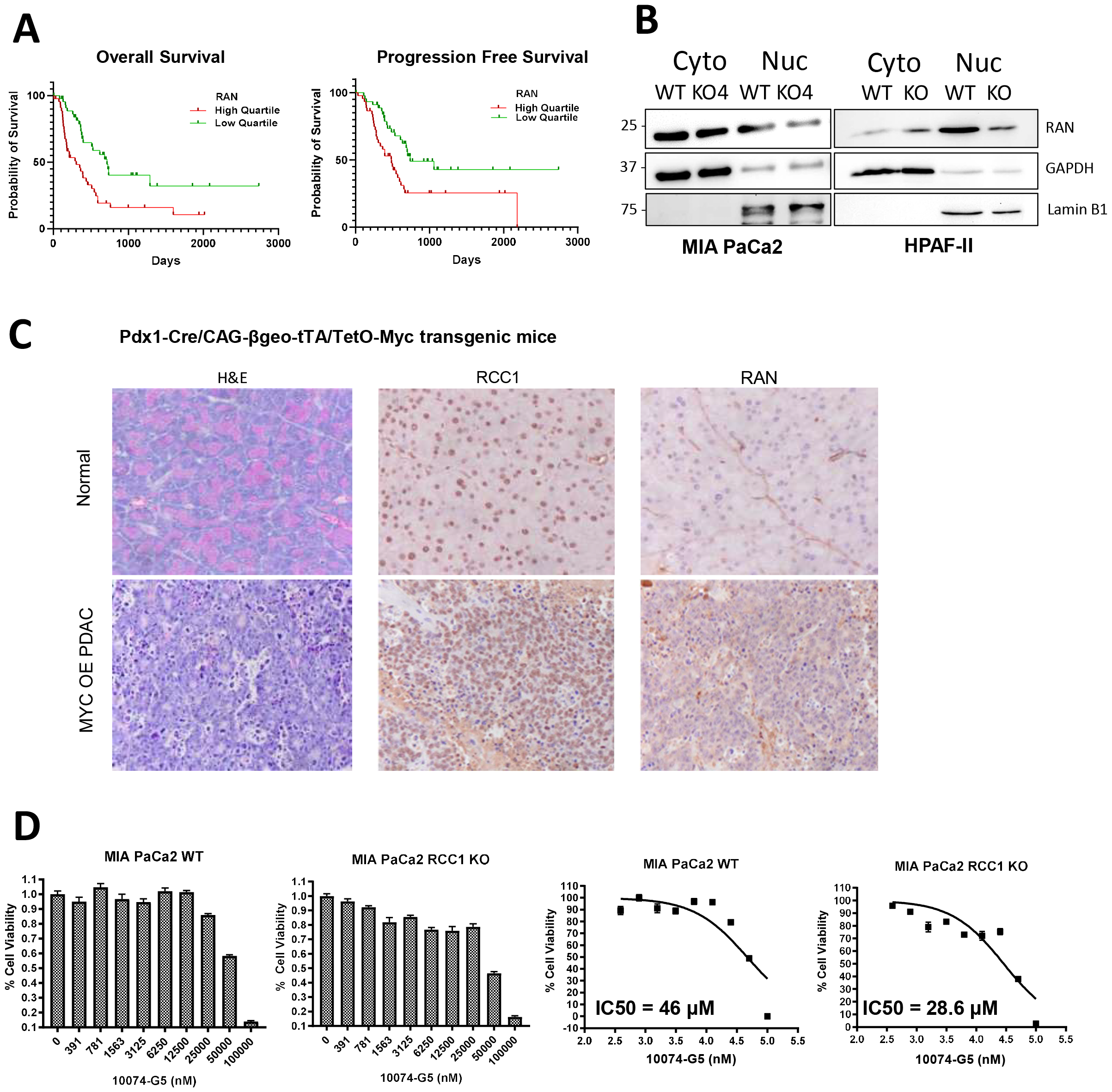
Myc-RCC1-Ran axis regulates PDAC. **(A)** Kaplan Meier plots of overall and progression free survival of PDAC patients in the TCGA in the high quartile vs the low quartile of *RAN* expression. **(B)** Nuclear cytoplasmic fractionation showing depleted nuclear Ran in MIA PaCa2 and HPAF-II *RCC1* KO cells. **(C)** H&Es and immunohistochemical staining of RCC1 and Ran in a c-myc overexpressing PDAC mouse model tissues. **(D)** MIA PaCa2 *RCC1* WT or KO cells were treated with gradually increasing concentrations of c-Myc inhibitor 10074-G5. IC50 values were calculated using non-linear regression analysis.

**Figure 7.**
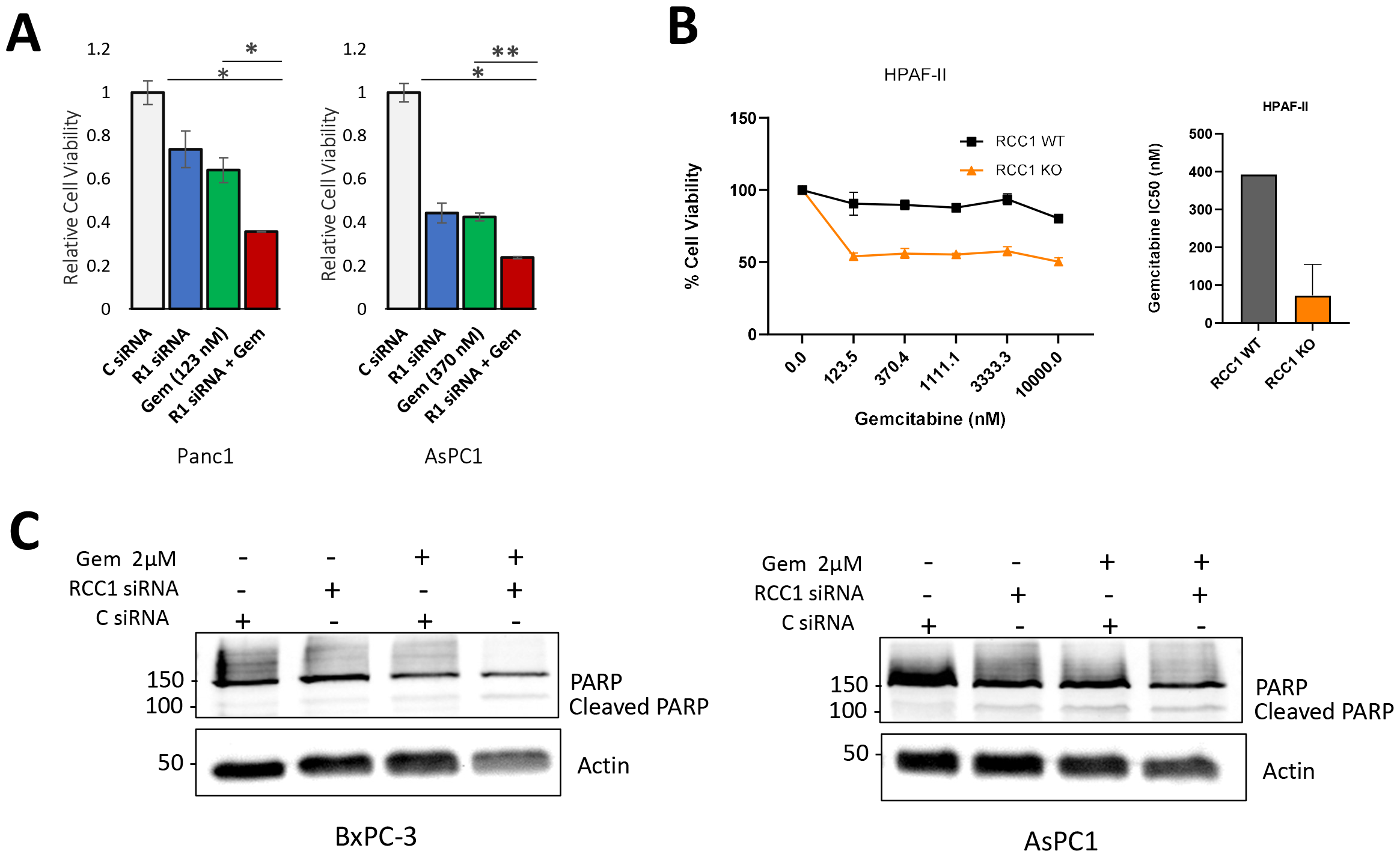
RCC1 depletion sensitizes cells to Gemcitabine treatment. **(A)** cells were transfected with Control or *RCC1* siRNA and treated with Gemcitabine. MTT was used to determine cell viability. siRCC1 in combination with Gemcitabine reduced cell viability. (B) HPAF-II cells with or without RCC1 were treated with increasing concentrations of Gemcitabine. Bar graph shows IC50 values. **(C)** Apoptosis analysis by cleaved PARP immunoblot in cells transfected with *RCC1* siRNA, treated with Gemcitabine, or a combination of both, showing enhanced apoptosis in the combination.

Analysis of alterations that correlate with higher RCC1 expression in patients revealed increased occurrence of c-Myc amplifications in patients with higher RCC1 status ^9^. We interrogated RCC1 and Ran levels in a c-Myc driven PDAC mouse model (Pdx1-Cre/CAG-βgeo-tTA/TetO-Myc) ^15^, where we found increased RCC1 expression in PDAC tissues (Figure 6C). Using a small molecule inhibitor of c-Myc, 10074-G5, which specifically inhibits the interaction between Myc and Max ^16^, we found that RCC1 KO cells are more sensitive to c-Myc inhibition compared to RCC1 WT MIA PaCa2 cells (Figure 6D). Additionally, we found that RCC1 KO cells have lower c-Myc levels, thus making the cells more sensitive to c-Myc inhibition (Figure S4D). These findings suggest that there may be a feedback loop that exists between RCC1 and c-Myc.

### RCC1 is involved in chemotherapy resistance and sensitization

There is some evidence to show that RCC1 is involved in the regulation of the DNA damage response (DDR) through transport of DDR factors in and out of the nucleus ^8^. Our data show that RCC1 KO results in redistribution of cell cycle phases within the cells, causing these cells to arrest in S and G2 phases (Figure 3G, S1D, S2D). We hypothesized that this increased replication stress in the cells, as well as the alteration in steady state nucleocytoplasmic transport may result in increased cell sensitivity to the standard of care chemotherapeutic agent Gemcitabine. PanC1 and AsPC1 cells showed enhanced sensitivity to Gemcitabine-mediated cell death when combined with RCC1 siRNA compared to gemcitabine alone (Figure 7A). HPAF-II cells with RCC1 KO also demonstrated increased sensitivity to Gemcitabine across a range of Gemcitabine concentrations and a reduction in IC50 of the drug compared to RCC1 WT cells (Figure 7B). Increased cleaved PARP in cells treated with Gemcitabine and RCC1 siRNA combination indicates that decreased cell viability upon treatment is at least partially due to increased apoptosis in these cells (Figure 7C). Overall, these data show that silencing of RCC1 results in enhanced sensitivity to Gemcitabine treatment.

## DISCUSSION

Cancer cells become addicted to Ran-mediated nuclear cytoplasmic transport for continuous growth, development, and progression. This creates an opportunity to target this cycle on which proliferative cells are more dependent compared to normal and quiescent cells. For the Ran GTPase to properly cycle between nuclear and cytoplasmic compartments, and between GTP-bound and GDP-bound states, two major players, RCC1 and RanGAP1 act on Ran. The localization of RCC1 and RanGAP1 is mutually exclusive allowing RCC1 to exclusively act on Ran in the nucleus generating high concentrations of RanGTP. On the other side of the nuclear membrane, RanGAP1 catalyzes RanGTP hydrolysis to RanGDP. In PDAC tissues, we found that elevated RCC1 levels on the RNA and protein levels. We also found that high RCC1 and high Ran expression are correlated with worse patient survival. This demonstrates that RCC1, which acts downstream of intractable signaling cascades, presents a novel targeting opportunity for PDAC.

Using siRNA transient and CRISPR-Cas9 stable silencing of RCC1, we examined the effects of RCC1 inhibition on PDAC. Cellular growth was inhibited, and xenograft tumor growth was markedly reduced. Mechanistically, we found that RCC1 is involved in the regulation of subcellular Ran GTPase distribution, and the maintenance of Ran concentrated within the nucleus. This leads to pervasive effects on the expression and localization of nuclear and cytoplasmic proteins as revealed by proteomic analysis. Identified altered targets within the proteomic analysis could provide opportunities for combination targeting.

The PDAC therapeutic landscape suffers from limited therapeutic efficiency of available drugs and a scarcity of targeted therapeutic options. The major oncogenic driver in PDAC is a mutant KRAS, which had been deemed undruggable until recently. The KRAS G12C mutation has been targeted with two drugs, adagrasib and sotorasib. However, KRAS G12C are uncommon in PDAC patients, and the new therapeutic agents may have limited therapeutic efficacy. Novel KRAS G12D inhibitors are in early clinical development, and results from these trials are eagerly awaited. Nonetheless, response to KRAS targeted therapies is usually limited primarily because of the rapid development of resistance. Therefore, the identification of novel drug targets and vulnerabilities for PDAC is essential to enhance treatment options for patients. RCC1 silencing in PDAC cells resulted in the sensitization to Gemcitabine treatment, which is a first-line chemotherapeutic agent for PDAC. Cells also showed an enhanced apoptotic response when RCC1 silencing was combined with chemotherapy. Overall, we have demonstrated that RCC1 is a promising therapeutic target for PDAC. Targeting RCC1 shows enhanced growth restriction in combination with Gemcitabine, which supports further investigation of this target.

## MATERIALS AND METHODS

### Cell culture

Human pancreatic ductal adenocarcinoma cells were maintained in appropriate media as follows: MIA PaCa2, PanC1, HPAC in DMEM (Gibco), BxPC3, ASPC1 in PRMI 1640 (Gibco), and HPAF-II in EMEM (Gibco). Media was supplemented with 10% FBS and 1% penicillin-streptomycin and cells were incubated in a 5% CO2 atmosphere at 37C. All cell lines have been authenticated in a core facility of the Applied Genomics Technology Center at Wayne State University, using short tandem repeat (STR) profiling on the PowerPlex 16 System (Promega). The cell lines were routinely tested for the absence of Mycoplasma species using PCR. Murine KCI313 cells were maintained in DMEM supplemented with 10% FBS and 1% penicillin-streptomycin.

### Human tissue

Rapid autopsy tissue collection was approved by the Wayne State University Institutional Review Board (IRB) # IRB-20-03-2022-M1 and Protocol No.: 2020-028. A written informed consent was obtained from the patients and the studies were conducted in accordance with Declaration of Helsinki and were approved by an IRB.

### Cell viability assays

Cells were seeded in 96-well microtiter culture plates at equal densities and allowed to grow for 72 hours. The reagent 3-(4,5-dimethylthiazol-2-yl)-2,5-diphenyltetrazolium bromide (MTT) was added to the wells at a final concentration of (0.5 mg/mL) and incubated for 3 hours at 37 degrees C. After the incubation period, the reaction was terminated by aspirating the supernatant and the addition of DMSO to solubilize the formazan crystals. After 15 minutes of gentle shaking, the absorbance was measured at 570 on SynergyHT (BioTek, Winooski, WI, USA) plate reader. Relative viability was calculated using absorbance data and plotted using GraphPad Prism 10.

### Transient knockdown and overexpression

Cells were seeded in a 6-well plate and incubated overnight to reach 90% confluence the next day. The cells were then transfected with the siRNA for knockdown or plasmid DNA for overexpression using lipofectamine 3000 (Life Technologies) according to manufacturer’s instructions, diluted in serum-free RPMI media. Plasmid-lipid complexes were added to the wells, fresh media was added 24 hours later. For knockdown assays, the cells were harvested 72 hours post-transfection for further assays. For proteomics analysis, the cells were collected and reseeded 48 hours after the first transfection and were transfected for the second time in a double transfection protocol prior to nuclear cytoplasmic fractionation. For Overexpression, cells were harvested 48 hours after transfection for RT-qPCR or re-seeded for further assays.

### Cell viability assay

Cells were seeded in 96 well plates at 3-4 × 10^3^ cells per well, then either transfected with siRNA or treated with drug the following day. After 48-72-hour incubation period, 3-(4,5-dimethylthiazol-2-yl)-2,5-diphenyltetrazolium bromide (MTT) reagent was added at a final concentration of 0.5 mg/ml. The plates were incubated for 3 hours to allow the live cells to form crystals of formazan. After the incubation period, the media was removed and 100 μL of DMSO were added per well. The plates were rocked gently for 15 minutes, and the absorbance was read using a SynergyHT plate reader at a wavelength of 570 nm. Viability was plotted using GraphPad Prism Software.

### Clonogenic assay

Cells were counted and reseeded at a density of 500 cells per well in a 6-well culture plate. Cells were left to grow for 14 days to form colonies. Plates were washed using phosphate buffered saline (PBS), fixed with methanol and stained with crystal violet for 30 minutes. The plates were then washed in tap water, then dried and photographed.

### Spheroid formation assay

A single suspension containing 1000 cells per 100 μL was plated in 96-well ultra-low attachment plates (corning) in 3D tumorsphere media XF (PromoCell). Media was added to the well every 3 days and growth was monitored microscopically. After spheroids were formed, they were photographed, and viability was quantified using cell titer glow assay (Promega).

### Nuclear cytoplasmic fractionation

Cells were fractionated into nuclear and cytoplasmic fractions using NE-PER Nuclear and Cytoplasmic Extraction Kit (Pierce Biotechnology), according to the manufacturer’s instructions. Briefly, trypsinized cells were washed twice with PBS by centrifugation at 500× g for 5 minutes. Ice-cold CER I was added to the cell pellet, mixed by vortexing, and CER II was added to the cells after 10 minutes of incubation. The tubes were vortexed and then centrifuged at maximum speed for 10 minutes at 4° C. The supernatant (cytoplasmic fraction) was transferred to a new tube. The pellet which contains the nuclei was then washed with ice-cold PBS, then incubated in ice-cold NER for 40 mins. The tubes were then centrifuged at maximum speed for 10 minutes to separate the nuclear extract. Protein concentrations of the fractions were measured, and the fractions were then used for subsequent Western blot analysis (as described below), or proteomics analysis.

### Scratch wound healing assay

Confluent cells in 6-well plates were scratched with the tip of a 200 μL pipette tip, then washed and replaced with fresh media. A line perpendicular to the scratch was drawn on the well to ascertain the images are taken in the same spot. Zero, 24, and 48-hour images were captured using an inverted microscope fitted with a camera.

### Immunoblot analysis

For total protein extraction, cultured cells were lysed in RIPA buffer (Santa Cruz Biotechnology) with protease and phosphatase inhibitors, and the protein concentration was measured using BCA protein assay (Pierce). Samples were prepared for electrophoresis in 6x Laemmli’s buffer and boiled for 5 minutes. The protein samples were separated by 10% SDS-PAGE, and subsequently transferred onto a nitrocellulose membrane. Membranes were blocked in 5% bovine serum albumin (BSA) in PBS containing 1% Tween (PBST) for one hour at room temperature prior to incubation in a specific primary antibody overnight at 4°C; specific conjugated secondary antibodies were added for 1 hour at room temperature (RT). The signal was detected by chemiluminescence (Pierce). ImageJ Software was used for densitometric quantification. The following antibodies were used: RCC1: Abcam, ab181155, 1:1000 dilution or Santa Cruz Biotechnology, sc-55559, 1:1000 dilution. Actin: Santa Cruz Biotechnology (SCBT), sc-8432, 1:2500 dilution. GAPDH SCBT: sc-365062 or sc-47724, 1:2500 dilution. Ran: Santa Cruz Biotechnology, or cell signaling #4462 1:1000 dilution. PARP #9542 1:1000 dilution, Lamin B1 SCBT: sc-377000 or cell signaling #13435 1:1000 dilution, HDAC3 cell signaling # 3949 1:1000 dilution, Histone H3.3 #9715 1:1000 dilution, Tubulin SCBT sc-73242 1:1000 dilution.

### RNA isolation and mRNA Real-Time qRT-PCR

Total RNA was purified from cell cultures using the RNeasy Plus Mini Kit (Qiagen), according to the manufacturer’s protocol. RNA was reverse transcribed using the High-Capacity cDNA Reverse Transcription Kit (Applied biosystems), and mRNA expression was analyzed by real-time qPCR using SYBR Green Master Mix (Applied Biosystems). The qPCR was initiated by 10 min at 95 °C followed by 40 thermal cycles of 15 s at 95 °C and 1 min at 60 °C in a StepOnePlus real-time PCR system (Applied Biosystems). Table 1 lists the sequences of the primers used.

### LC/MS-MS proteomics

Proteins were extracted using nuclear cytoplasmic fractionation as described above. Proteins were quantified using label-free quantitative proteomics after digestion. A total of 3113 proteins were identified. To identify differentially expressed proteins, the fold-change cutoff was set at 1.2. To understand the functions of identified proteins, KEGG pathway and GO annotation and enrichment analyses were performed.

### Whole exome RNA sequencing and bioinformatics analysis

Messenger RNA was purified by poly-T oligo attached magnetic beads from total RNA. After fragmentation, the first strand cDNA was synthesized using random hexamer primers, followed by the second strand cDNA synthesis using either dUTP for directional library or dTTP for non-directional library. For the non-directional library, it was ready after end repair, A-tailing, adapter ligation, size selection, amplification, and purification. For the directional library, it was ready after end repair, A-tailing, adapter ligation, size selection, USER enzyme digestion, amplification, and purification. The library was checked with Qubit and real-time PCR for quantification and bioanalyzer for size distribution detection. Quantified libraries were pooled and sequenced on Illumina platforms, according to effective library concentration and data amount. Gene expression level was estimated by the abundance of transcripts that mapped to the human reference genome, estimated as Fragments Per Kilobase of transcript sequence per Millions base pairs sequenced (FPKM). Differential gene expression was analyzed using DESeq2 ^17^.

### Immunofluorescence imaging

Cells cultured on chambered glass slides were fixed with 4% paraformaldehyde for 10 min and subsequently permeabilized with 0.05% Triton-X100 for 15 min. The slide chambers were then washed with PBS and blocked in 5% BSA solution for one hour. Slide chambers were washed again with PBS, and a specific primary antibody (1:100 dilution in 3% BSA) was added and incubated overnight with gentle rocking at 4 °C. The next day, slides were washed with PBS-T and PBS before goat anti-rabbit secondary antibody Alexa Fluor® 488 conjugate (Invitrogen) was added at 1:500 dilution in 3% BSA. Finally, Slides were mounted with ProLong™ Gold antifade reagent with DAPI (Life Technologies) and a coverslip. An inverted fluorescent microscope was used for imaging.

### Flow cytometric apoptosis analysis

Cells were sorted using Annexin V FITC and 7-amino-actinomycin D (7-AAD) according to the manufacturer’s protocol. Cells were seeded in a 6-well plate and transfected with siRNA for 48hours prior to trypsinization and staining with Annexin V and PI, or 7-AAD. Stained cells were sorted using the Becton Dickinson FACSCanto or the Cytek Northern Light flow cytometer at the Karmanos Cancer Institute Microscopy, Imaging and Cytometry Resources (MICR) Core.

### Cell cycle assay

Cell cycle phase was determined using DNA content analysis by flow cytometry. Cells were harvested and fixed in ice cold 70% ethanol, and left overnight at 4°C. The following day, the cells were washed in PBS, then permeabilized with Triton and stained using PI with DNase free RNase. DNA content was detected using Becton Dickinson FACSCanto or the Cytek Northern Light flow cytometer at the Karmanos Cancer Institute Microscopy, Imaging and Cytometry Resources (MICR) Core.

### Mouse xenograft model of PDAC

All studies were conducted under Wayne State University’s Institutional Animal Care and Use Committee approved protocol. One million cells of HPAF-II or MIA PaCa2 cell lines (RCC1 WT or KO) were injected subcutaneously into 6-week-old ICR-SCID mice. Cells were suspended in 100 μL PBS, loaded in BD 26G 1 mL sub-Q syringe and injected bilaterally in the flanks of mice. Tumor sizes were followed by caliper measurements.

### Immunohistochemistry (IHC)

IHC was performed on formalin fixed paraffin embedded (FFPE) sections mounted on super frost slides. Paraffin sections were de-waxed in a xylene-ethanol series. Endogenous peroxides were blocked by a methanol/3% hydrogen peroxide incubation at room temperature for 30 minutes. HIER was performed with a pH 6 citrate buffer and pH 9 buffer, using the BIOCARE Decloaking Chamber. A 40-minute blocking step with Super Block Blocking buffer (Thermo Scientific) was performed prior to adding the primary antibody. Detection was obtained using GBI Labs DAB chromogen kits and counterstained with Mayer’s Hematoxylin. Sections were then de-hydrated through a series of ethanol to xylene washes and cover slipped with Permount.

Mouse xenograft tissues were processed and stained with H&E stains or IHC using RAN antibody (Cell Signaling, 4462S, dilution of 1:100, overnight at 4°C) RAP and TMA sections IHC stained with anti-RCC1 antibody (RCC1, Abcam, ab181155, dilution of 1:100, overnight at 4°C). This work was done at the Pathology Core facility at Wayne State University.

### Statistical analysis

All experiments were performed in triplicates unless otherwise noted. (*, P < 0.05; **, P < 0.01, *** P < 0.001) were determined by two-sided unpaired t tests, unless otherwise noted.

### Data availability

The data generated in this study are made available upon request from the corresponding author.

## Acknowledgements

Karmanos’ support for the Pancreatic Tumor Organ Donation Program is Acknowledged. SFB acknowledges support from the DeRoy Testamentary Foundation.

