## Supplementary figures and images for "Regulator of Chromosome Condensation (RCC1) a novel therapeutic target in pancreatic ductal adenocarcinoma drives tumor progression via the c-Myc-RCC1-Ran axis"

### Supplemental Figures

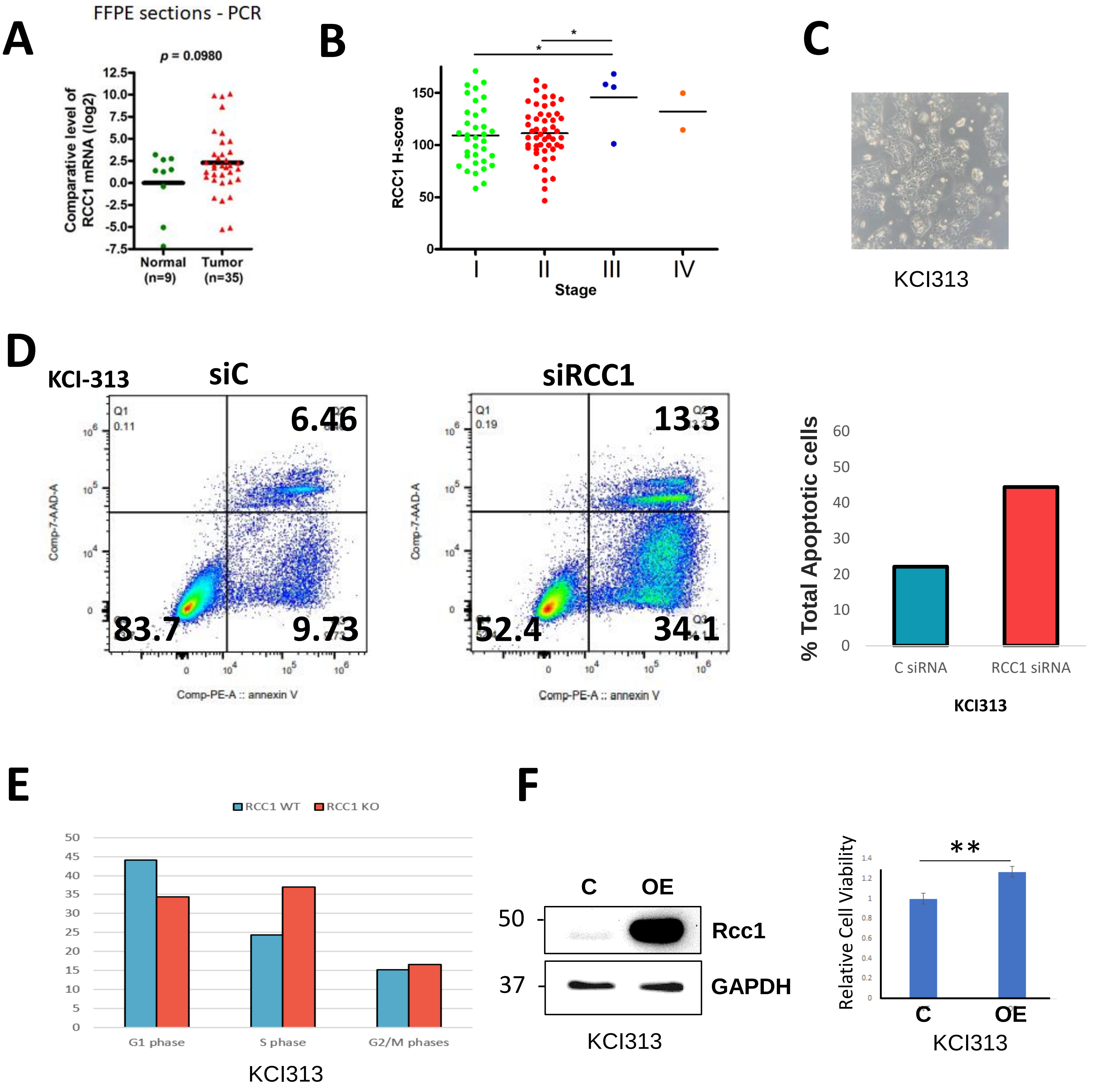

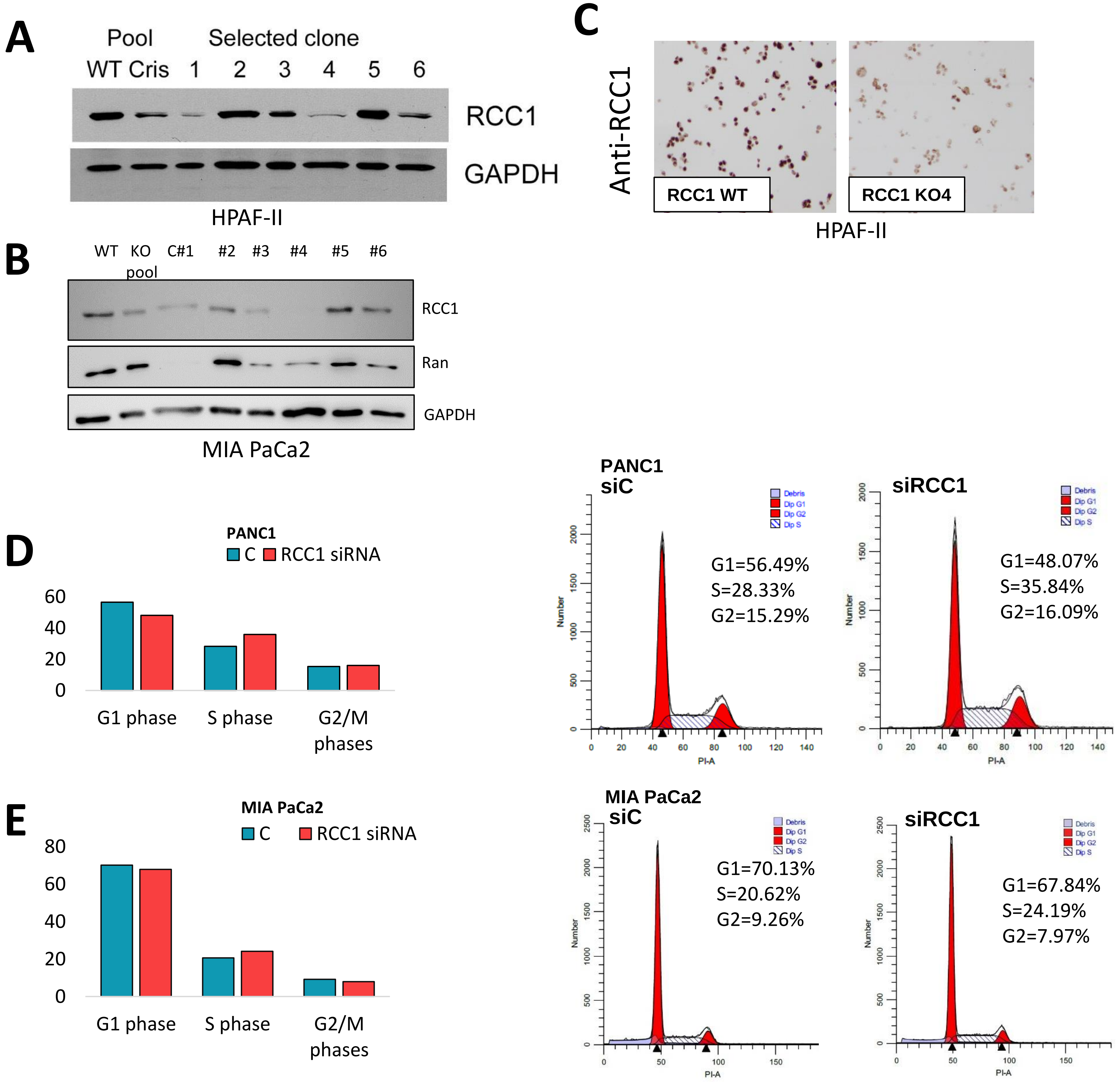

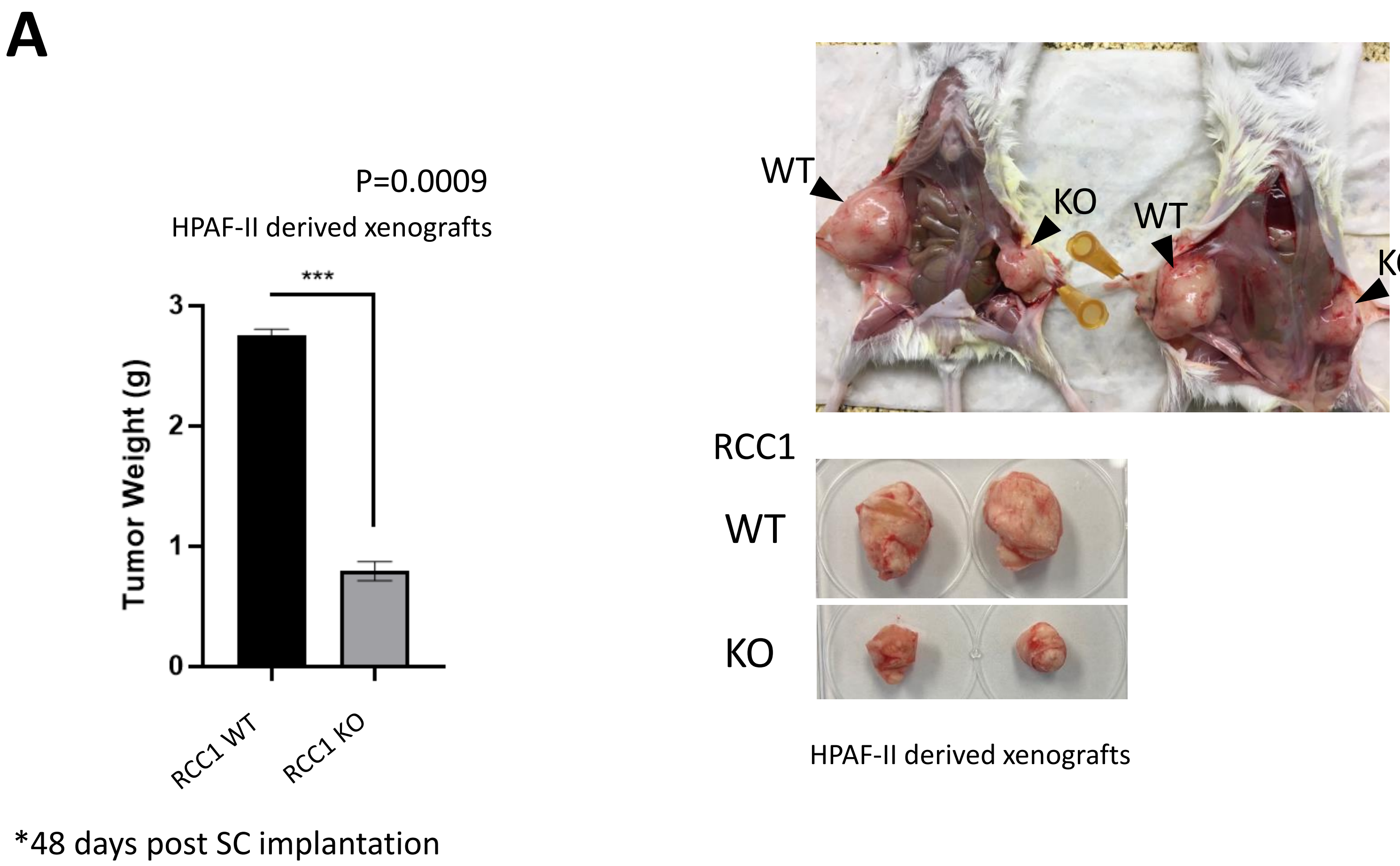

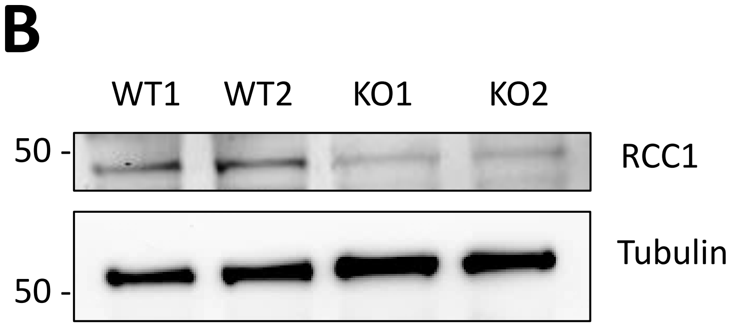

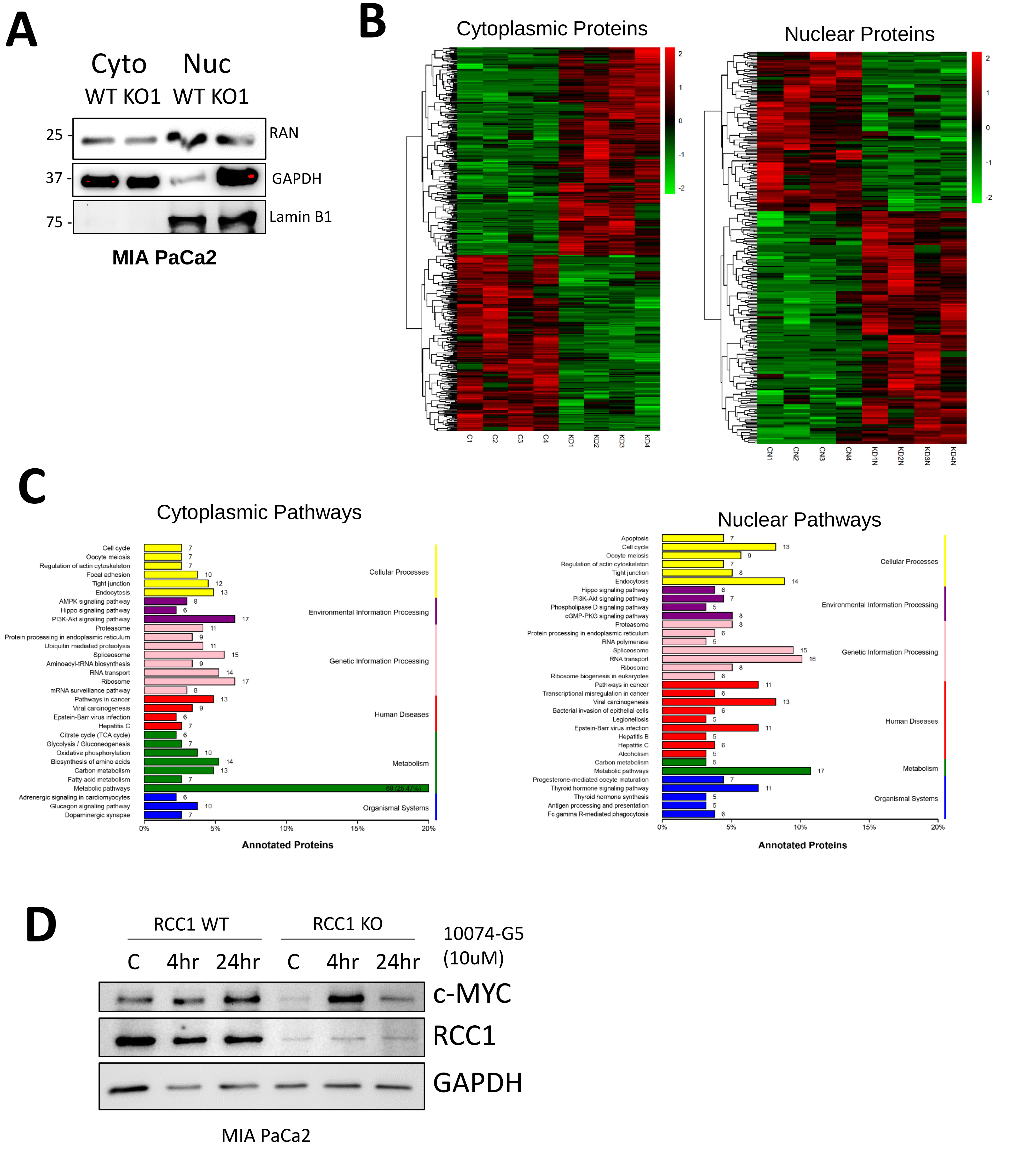
